# Bursty Expression Enhances Reliability of Messaging in Gene Networks

**DOI:** 10.1101/2020.02.09.940684

**Authors:** Leonardo R. Gama, Guilherme Giovanini, Gábor Balázsi, Alexandre F. Ramos

## Abstract

The promoter state of a gene and its expression levels are modulated by the amounts of transcription factors interacting with its regulatory regions. Hence, one may interpret a gene network as a communicating system in which the state of the promoter of a gene (the *source*) is communicated by the amounts of transcription factors that it expresses (the *message*) to modulate the state of the promoter and expression levels of another gene (the *receptor*). The reliability of the gene network dynamics can be quantified by the Shannon’s entropy of the message and the mutual information between the message and the promoter state. Here we consider a stochastic model for a binary gene and use its exact steady state solutions to calculate the entropy and mutual information. We show that a slow switching promoter having long and equally standing ON and OFF states maximizes the mutual information and reduces entropy. That is a bursty regime generating a high variance message governed by a bimodal probability distribution with peaks of the same height. Our results indicate that Shannon’s theory can be a powerful framework for understanding how bursty gene expression conciliates with the striking spatio-temporal precision exhibited in pattern formation of developing organisms.

## 1 Introduction

The remarkable spatio-temporal precision observed during development of *metazoa* is the result of well orchestrated gene networks regulating the expression of proteins that determine cell fate. That contrasts with the unavoidable fluctuations of intracellular molecular numbers caused by low copy numbers of reactants [1] as detected in expression of prokaryotic and eukaryotic cell’s genes [2–17]. Hence, one may ask whether in the due course of evolution there emerged mechanisms equipping biological systems with the capacity of modulating noise to exploit it functionally [18]. One possible example is bursty gene expression which appears to be an ubiquitous transcriptional regime occurring in developing *Drosophila* embryos [19–21]. During development, pattern formation is regulated by gene networks generating protein levels within proper ranges at proper instants and positions of the embryo [11, 19–25]. For those patterns to be achieved, the activation and inactivation of the promoters of the genes of those networks are modulated with sufficient precision. Hence, one may ask how the promoters, and transcription factors regulating the expression of these gene networks, contribute to the emergence of such a reliable dynamics.

Here we approach those questions considering a gene network as a communicating system in which a gene (the source) expresses transcription factors (the message) that regulate another gene (the receptor). That enables the application of information theory for investigating transduction by a gene network of input signal into functional output [26–28]. As opposed to the common consideration of gene activation/inactivation as completely random events, here we assign to them information content originating, for example, from upstream regulators. Therefore, we assume the information content of the input message to be the promoter state of the source, since the binding of transcription factors to the enhancer induces the state of the promoter of the receptor gene [29, 30]. The processing of all information arriving at the receptor is not considered here [31–34], as we are interested on the reliability of the information content of the message. Hence, we employ Shannon’s theory to compute the entropy of the message and the mutual information between the message and the promoter state of the source [35] by means of the probabilities given by the steady state exact solutions for the stochastic model for binary gene expression [36–38]. Since we are analysing the information content of the message we may interpret the number of products expressed by the binary gene as mRNA’s (which would result on the transcription model) or as transcription factors (which would result on a coupled transcription-translation model). Our results show that the gene having a slow switching promoter with ON and OFF states of the same duration maximizes mutual information between the message and the promoter state while reducing the message’s entropy. That reveals a new perspective to understand the origins of the precision observed in biological systems based on Shannon’s framework as a useful tool for investigating its mechanisms.

### 1.1 Preliminaries

Fig. 1 shows the effective set of reactions used to interpret a gene network as a communicating system. It depicts the symbols for: synthesis rate of the transcription factors when the promoter is ON, *k*; transcription factors degradation rate, *ρ*; promoter switching rate from ON to OFF state and vice-versa, *h* and *f*, respectively; the number of transcription factors available, *n*. No synthesis happens when the promoter is OFF. The curved lines indicate the interaction of the protein encoded in one gene with the enhancer of the receptor.

**Figure 1:**
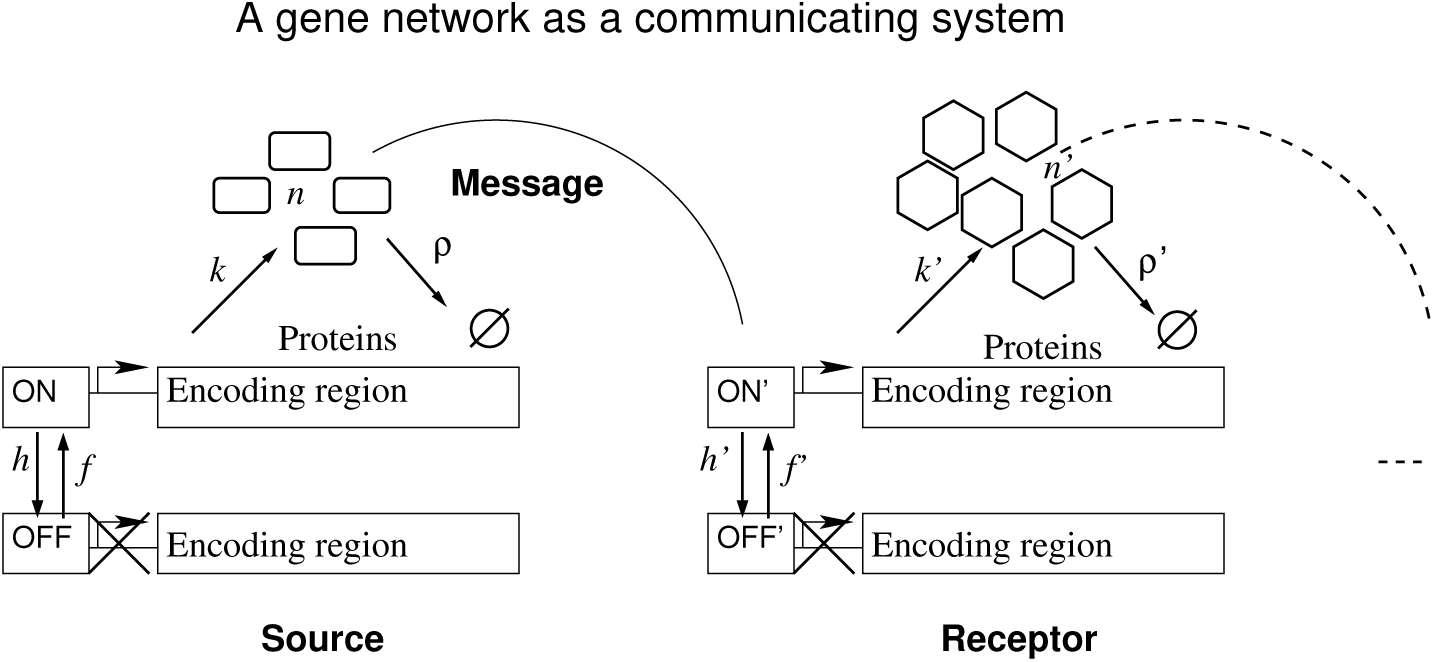
Two elements of an arbitrary gene network are depicted as a communicating system. The continuation of the network is represented by the dashed lines. A source gene communicates with a receptor gene through transcription factors and the information is the promoter state of the source.

The average number of transcription factors (or expected message from the source) is denoted as usual by ⟨*n*⟩. The bursting (hyper-Poissonian) nature of the message can be measured by the Fano factor, ℱ, defined as the ratio of the variance to the mean (see Eq. 1). A Poissonian distribution has ℱ = 1 and a super Poissonian distribution has ℱ = 1 + *δ*, where the increment *δ* is positive [2, 39]. We denote as bursty the expression of a gene for which its products are governed by super Poissonian distributions. In some cases, bursty expression can generate bimodal distributions, with numbers of protein products governed by distributions having two distinct peaks. The bimodal distributions are consistent with trajectories of the number of gene products characterized by a repetition of the following events, assuming that initially the promoter is OFF and there are no transcription factors : a burst towards a finite value ⟨*n*_*α*_⟩ after the promoter switching ON; small fluctuations around ⟨*n*_*α*_⟩ while the promoter remains ON; a sharp reduction towards zero after the promoter switching OFF; small fluctuations around zero while the promoter remains OFF; and the cycle repeats.

The information content of the message is quantified in the framework of Shannon’s communication theory, where the variance of binary signals correlates with their information content. Hence, the average information content of the message is given by Shannon’s entropy; the coupling between the message and the state of the promoter of the source is given by the mutual information; the average information content of the message given the promoter being ON is measured by the conditional entropy (see Subsec. 4.7). We assume that the stochastic binary gene expression model is at the steady state limit. Hence, the promoter states and the number of transcription factors will change accordingly with constant probabilities. The fraction of durations of the ON and OFF states of the promoter is governed by constant marginal probabilities *p*_*α*_ and *p*_*β*_, respectively (see Eq. 7), for the promoter to be ON and OFF. For the promoter being exclusively ON the source would become constitutive (see Subsec. 4.1) with average number of transcription factors being *N* (see Eq. 3 or Eq. 7). The probability of finding *n* transcription factors and the promoter ON is denoted by *α*_*n*_ while *ϕ*_*n*_ indicates the marginal probability of finding *n* transcription factors (see Eq. 4 and Eq. 6).

The parameter *ϵ* is proportional to the ratio of the transcription factor’s lifetime to the average period of a complete cycle of promoter switching (see Eq. 7) for a fixed *p*_*α*_. For *ϵ* ≪ 1 the switching has the longer time scale and enables the average number of transcription factors to reach steady state value before the promoter switches from one state to another. For *ϵ* ≫ 1 we have the opposite condition and there will be multiple promoter switchings during the lifetime of a transcription factor.

## 2 Results

### 2.1 Bursty expression enables entropy reduction and mutual information increase

The horizontal axis of both graphs of Fig. 2 shows the bursting quantified by the Fano factor (see Eq. 1). The vertical axis of graphs A and B shows, respectively, the entropy of the message (see Eq. 11) and the mutual information between the message and the state of the promoter of the source (see Eq. 14). Each continuous line is associated to a specific value of the promoter switching rate, *ϵ* (see Eq. 7). The dashed curves are isolines for fixed values of the probability for the promoter to be ON, *p*_*α*_. The color scheme for the curves is given at the caption. The black solid diamond gives the entropy of the message when it is sent by a Poissonian source which is usually employed on the description of a constitutive gene. Note that we have no mutual information for the constitutive gene because we assume that its promoter will be always ON. A condition of equivalence between the binary gene to the constitutive one can be devised for *ϵ* >> *N* which corresponds to a switching cycle being much faster than the protein synthesis rate. In that case, the mutual information approaches zero.

**Figure 2:**
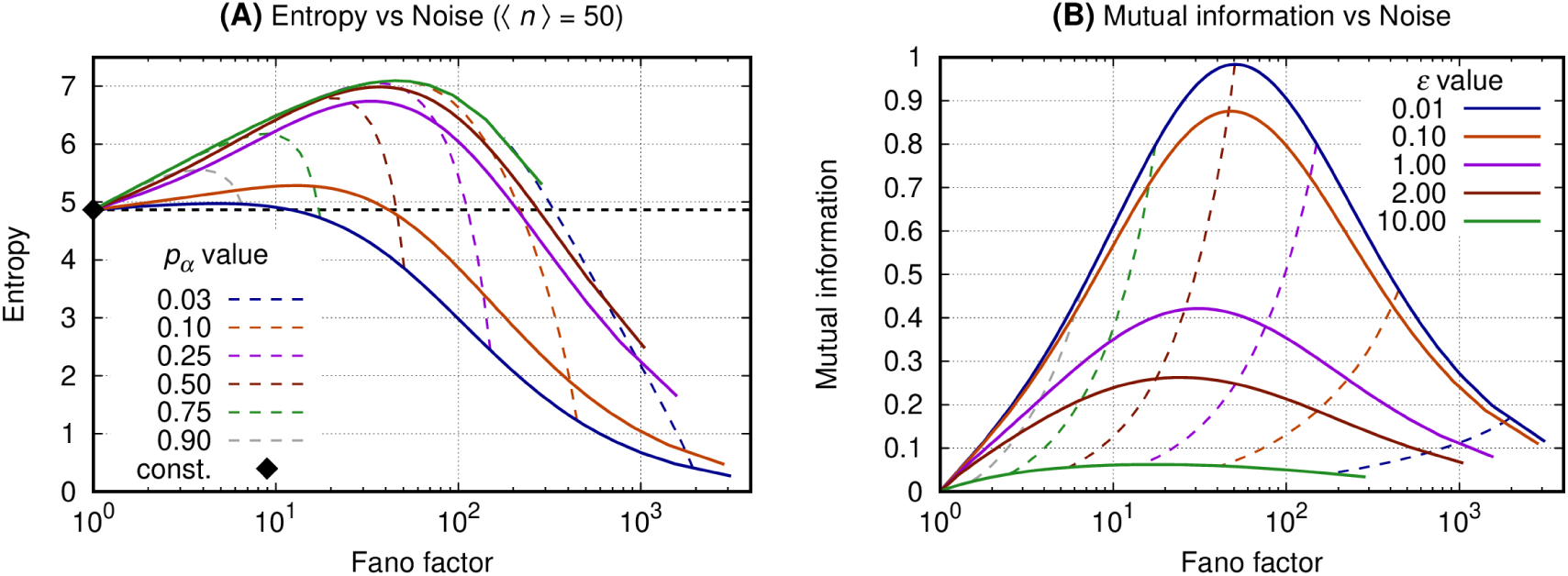
Entropy (or mutual information) and Fano factor were obtained varying *N* for fixed *p*_*α*_ and *ϵ*. All curves have the same value for the average number of gene products (⟨*n*⟩ = 50). Graph A shows the entropy of the message sent by the gene versus the noise of the message. Graph B shows the mutual information between the message and the source state as a function of the bursting. Continuous lines refer to fixed values of *ϵ* and fixed values of *p*_*α*_ are indicated by the dashed ones.

*Graph A* shows that the constitutive gene sets a reference value for the entropy and the binary source at the small bursting regime (ℱ ∼ 1) generating a message for which the entropy is greater than that sent by the constitutive gene. As bursting increases, however, the entropy of the message sent by the binary source reaches a maximum and then starts decreasing. Then the entropy of the message sent by the binary source becomes smaller than that of the constitutive source and decays towards arbitrarily small values. Besides, for a fixed value of ℱ, the message with minimal entropy is generated by a process having the smallest *ϵ*.

*Graph B* shows that the mutual information between the message and the source state reaches a maximum for values of the Fano factor being ∼ 1 + ⟨*n*⟩ and *ϵ* ≪ 1. For a fixed value of ℱ the mutual information becomes minimal for the greatest values of *ϵ*. As the switching rate becomes slower, *i*.*e. ϵ* ≪ 1, the mutual information is maximal for *p*_*α*_ ∼ 1*/*2. Furthermore, one may set an interval at which the mutual information has higher, significant values, by arbitrarily fixing the probability for the promoter to be ON within a range 1*/L* ≤ *p*_*α*_ ≤ 1 − 1*/L* such that the corresponding noise is (*L* − 1)⟨*n*⟩ ≥ ℱ ≥ ⟨*n*⟩*/*(*L* − 1), where *L* > 1 and ℱ ≫ 1 such that (ℱ − 1)(1 + *ϵ*) ∼ ℱ.

### 2.2 The slow switching genes generate reduced entropy and increased values for mutual information

The horizontal axis of both graphs of Fig. 3 indicates the values of the transcription factor’s lifetime relative to the ON and OFF switching cycle of the source’s promoter. The vertical axis of graphs A and B shows, respectively, the entropy of the message (see Eq. 11) and the mutual information between the message and the source state. To each continuous line we associate a specific value of the probability for the source to be ON, *p*_*α*_ (see Eq. 7). The dashed black curve indicates the entropy for the constitutive gene. This value is given as a reference since there is no switching on a constitutive promoter and, additionally, there is no mutual information between the promoter state and the message. The color scheme for the curves is given at the caption.

**Figure 3:**
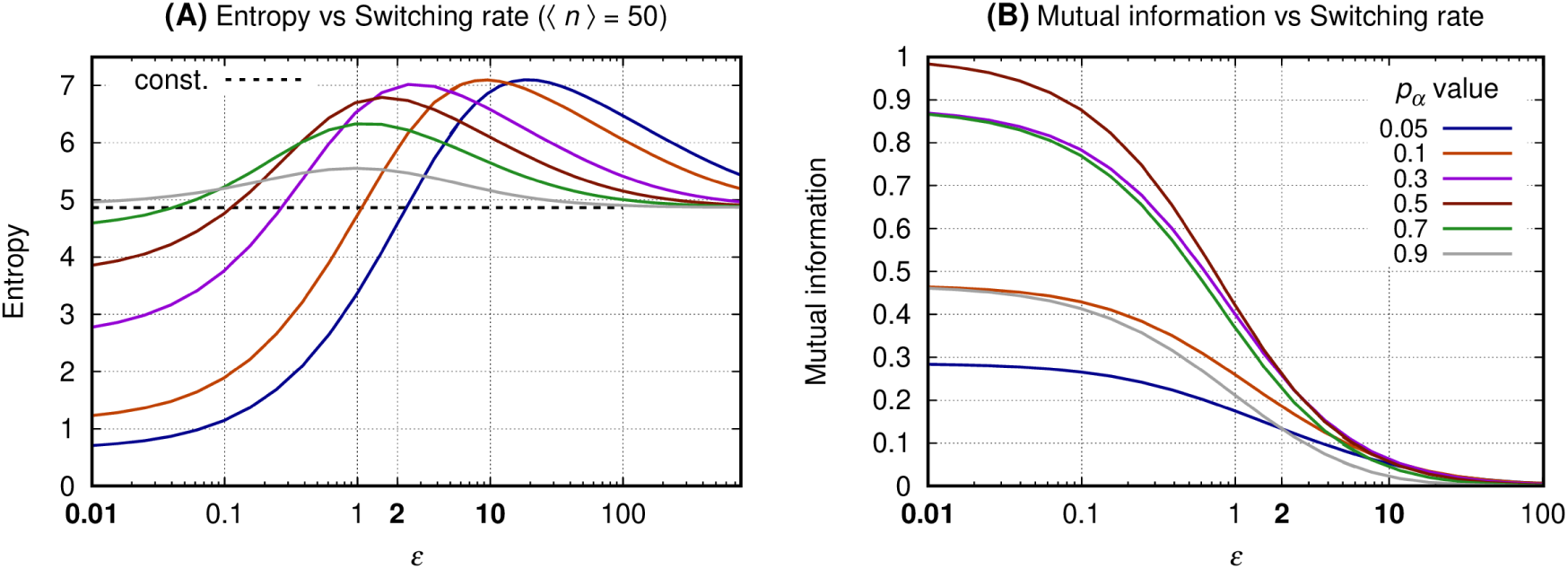
The entropy (and mutual information) was obtained fixing *p*_*α*_ and *N*. All curves have the same average number of ttranscription factors (⟨*n*⟩ = 50). The horizontal axis of both graphs indicate the transcription factor’s lifetime relative to the source’s ON and OFF promoter switching duration. Graph A shows the entropy of the message and graph B shows the mutual information between the message and the source’s promoter state. Each continuous line corresponds to a given value of *p*_*α*_ and the entropy value for a constitutive gene producing ⟨*n*⟩ = 50 proteins is shown by the horizontal dashed black line. Values of *ϵ* used to construct Fig. 4, Fig. 5, and Fig. 6 appear in bold at the horizontal axis label.

*Graph A* shows that the entropy for a given *p*_*α*_ is minimal when the switching is slow. As the switching rate increases, the entropy reaches a maximum value (*H*_*m*_) for an *ϵ*_*m*_ which is greater as the probability for the promoter to be ON is reduced. For *ϵ* > *ϵ*_*m*_ and a given *p*_*α*_, the entropy of the message decreases and asymptotically approaches that of a constitutive source. A source having *ϵ* ≪ 1 generates a message for which the entropy increases with *p*_*α*_ while the opposite happens for large values of *ϵ*. Furthermore, for *ϵ* ≪ 1 and *p*_*α*_ = 1*/*2 the entropy is smaller than that of the constitutive source, but not minimal.

*Graph B* shows the mutual information as a function of *ϵ* with each line being related to a specific value of *p*_*α*_. The similarity of the mutual information for complementary values of *p*_*α*_ as the switching rate increases relates to the bell-shaped curves (in semi-log scale) shown on Fig. 2 (graph B). For a given value of *p*_*α*_ the mutual information decreases monotonically with the increase of *ϵ*. For *ϵ* ≪ 1 the mutual information is maximal when *p*_*α*_ = 1*/*2, and for *p*_*α*_ ≪ 1 the message will have minimal entropy and minimal mutual information with the promoter state.

### 2.3 The distributions of the slow switching bursty regime with reduced entropy and maximal mutual information are bimodal

Figures 4–6 have four graphs each where: graph A shows the entropy on vertical axis and the average number of gene products (⟨*n*⟩) on the horizontal; graph B shows the entropy conditional to the promoter being ON versus ⟨*n*⟩_*α*_ (see Eq. 8); graph C shows the mutual information versus ⟨*n*⟩; graph D shows the probability distribution of *n*. Each continuous line correspond to a given *p*_*α*_ with color code given at the key of *graph B*. The values of the entropy (vertical axis of *graph A*); entropy conditional to the source being ON (vertical axis of *graph B*); mutual information (vertical axis of *graph C*) are obtained fixing a given *p*_*α*_ and varying *N* to generate the values of ⟨*n*⟩ at the horizontal axis. The probability distributions of *graph D* are obtained by also fixing ⟨*n*⟩ = 50, such that *N* = 50*/p*_*α*_. For each *p*_*α*_ we compute the values of *ϕ*_*n*_ (continuous lines) and *α*_*n*_(dashed lines). In each set of graphs the value of *ϵ* is fixed and given at graph A’s title. Results for the constitutive gene are shown by a black line on graphs A, B, and D.

**Figure 4:**
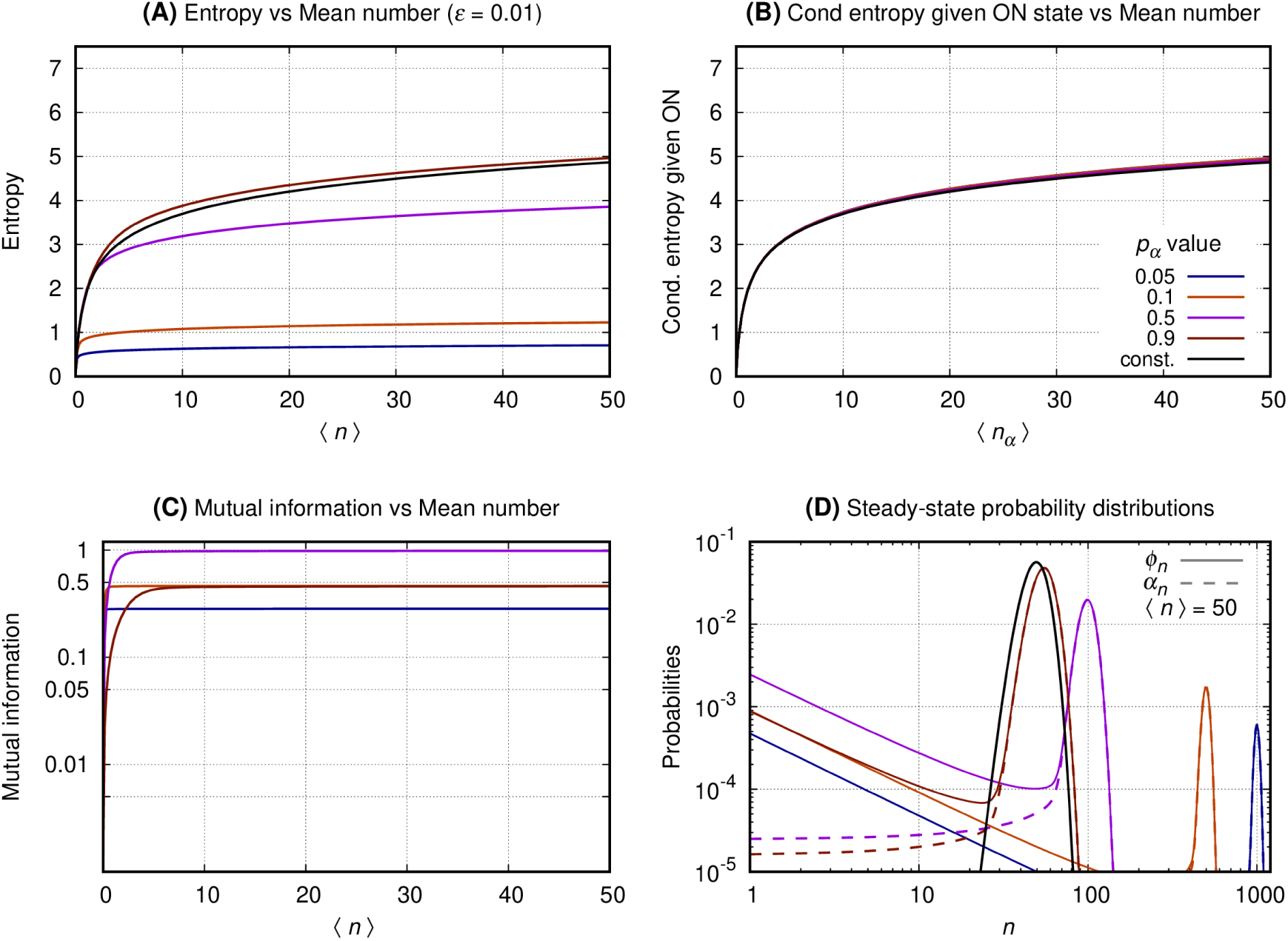
The switching rate of the promoter of the source is 0.01. The vertical axis of graph’s A, B, and C shows, respectively, entropy versus ⟨*n*⟩, entropy conditional to the source being ON versus ⟨*n*_*α*_⟩, and the mutual information versus ⟨*n*⟩. Graph D shows the probability distribution of *n*.

**Figure 5:**
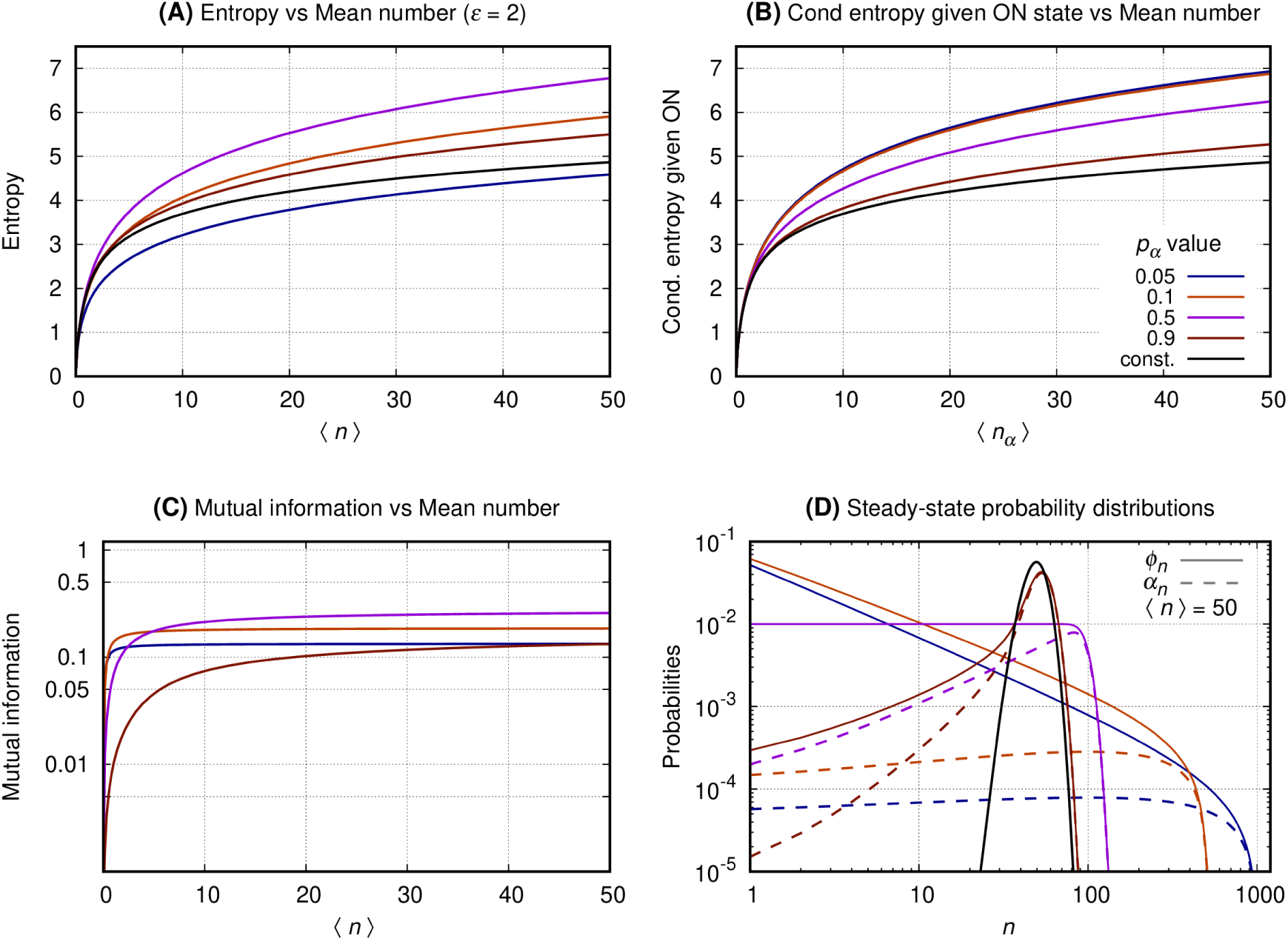
The switching rate of the promoter of the source is 2. The vertical axis of graph’s A, B, and C shows, respectively, entropy versus ⟨*n*⟩, entropy conditional to the source being ON versus ⟨*n*_*α*_⟩, and the mutual information versus ⟨*n*⟩. Graph D shows the probability distribution of *n*.

**Figure 6:**
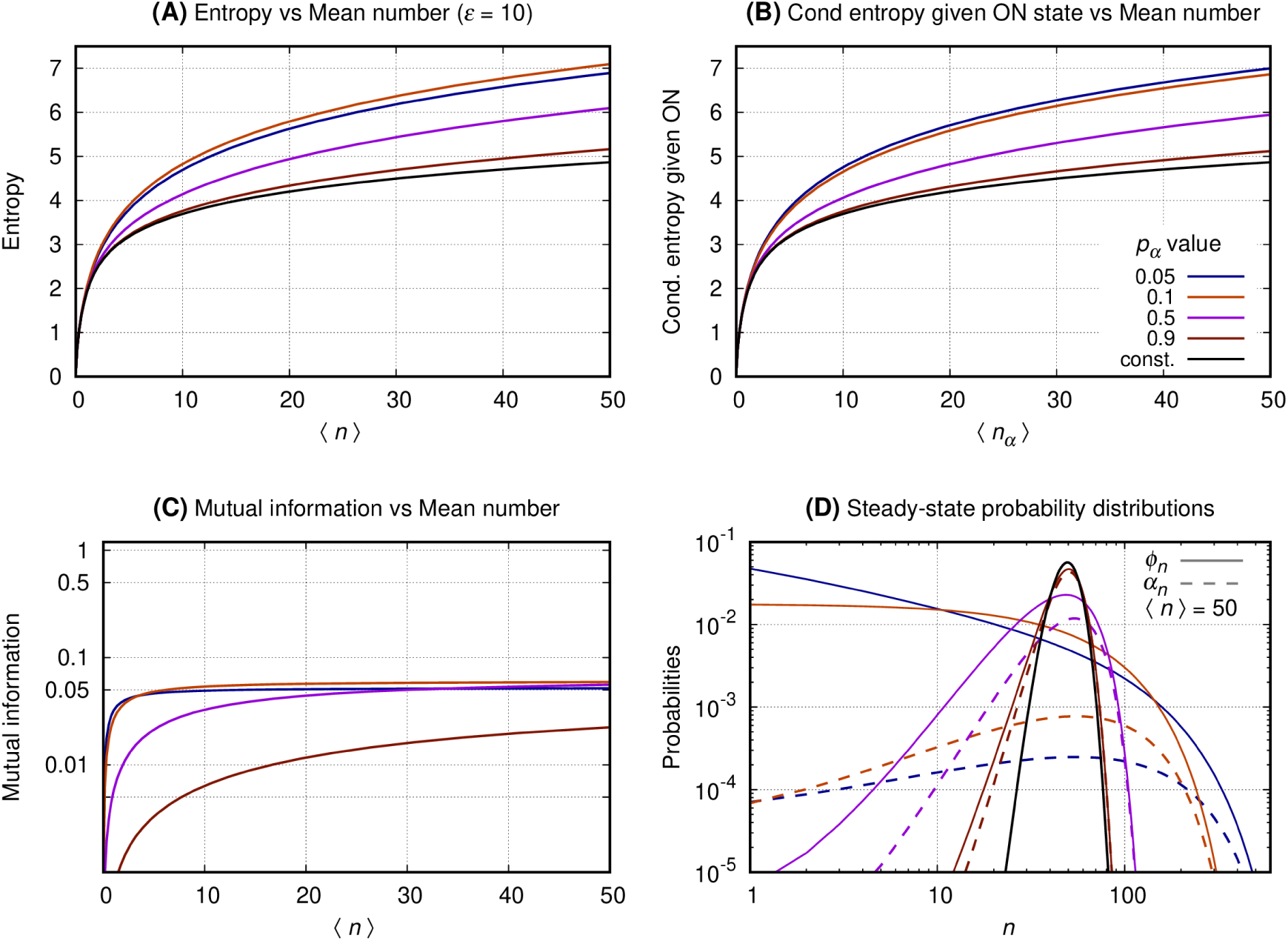
The switching rate of the promoter of the source is 10. The vertical axis of graph’s A, B, and C shows, respectively, entropy versus ⟨*n*⟩, entropy conditional to the source being ON versus ⟨*n*_*α*_⟩, and the mutual information versus ⟨*n*⟩. Graph D shows the probability distribution of *n*.

**Fig. 4**. *Graph A* shows that the entropy of the message increases monotonically with ⟨*n*⟩ for both the constitutive and binary sources. As *p*_*α*_ approaches one the entropy of the binary sources approaches that of a constitutive. A source having smaller values of *p*_*α*_ will send messages with significantly reduced entropy which growth with ⟨*n*⟩ is much slower. *Graph B* shows the entropy of the message under the condition of the source being ON. In that case the conditional entropy of all messages approach that of a constitutive source for all values of *p*_*α*_. That is observed by introducing the entropy for the constitutive source as a reference value (black curve). *Graph C* shows that the mutual information between the source’s promoter state and the message increases monotonically until it reaches a limit at a value that depends on *p*_*α*_. The mutual information for two complementary values of *p*_*α*_ approximate each other for a sufficiently large value of ⟨*n*⟩. For *p*_*α*_ = 1*/*2 the mutual information reaches its maximum with its plateau being reached when ⟨*n*⟩ ∼ 30. *Graph D* shows the probability distribution for ⟨*n*⟩ = 50. The black curve shows the probability distribution of *n* when emitted by a constitutive source. The distributions generated by the binary source are all bimodal, with one mode centered around zero and the other around *N* (where *N* = ⟨*n*⟩ */p*_*α*_). As the value of *p*_*α*_ approaches zero the second peak has its height lowered and its center displaced to the right. The opposite happens when *p*_*α*_ approximates unity with the taller peak approaching that of the constitutive source for the same mean.

**Fig. 5**. *Graph A* shows that the entropy of the message increases monotonically with ⟨*n*⟩ for both the constitutive and the binary sources. For larger values of *p*_*α*_, the entropy of the binary sources are greater than that of a constitutive. Entropy of the message from the binary source is a bit smaller than that of a constitutive for small values of *p*_*α*_. *Graph B* shows the entropy of the message given the promoter of the source being ON. In that case the conditional entropy of all messages is greater than that of a constitutive source. *Graph C* shows that the mutual information also increases monotonically until it reaches a limit value that depends on *p*_*α*_. The mutual information for two complementary values of *p*_*α*_ approximate each other for a sufficiently large value of ⟨*n*⟩. For *p*_*α*_ = 1*/*2 the mutual information reaches its maximum with its plateau being reached when ⟨*n*⟩ ∼ 30. Note, however, that the maximal value for the mutual information is smaller than that for *ϵ* = 0.01. *Graph D* shows the probability distributions for the binary source. The distributions generated by the binary source have significant probability values spread along a large range of values of *n*. For *p*_*α*_ = 1*/*2 the distribution is table shaped. As *p*_*α*_ → 1 the distribution forms a peak that gets closer to that of the distribution of a constitutive source while the distributions have a sector governed by a power law when *p*_*α*_ approaches zero.

**Fig. 6**. *Graph A* shows that the entropy of the message increases monotonically with ⟨*n*⟩ for both the constitutive and the binary sources. The entropy of the message emitted by the binary source is larger than that of a constitutive gene for all values of *p*_*α*_. *Graph B* shows the entropy of the message given the promoter being ON. The conditional entropy of all messages are greater than that of a constitutive source. *Graph C* shows that the mutual information also increases monotonically until it reaches a limit value that depends on *p*_*α*_ and that this maximal value is smaller than that for *ϵ* = 0.01 or *ϵ* = 2. *Graph D* shows the probability distributions for the binary and the constitutive sources. The distributions generated by the binary source are spread along a large range of values of *n*. The distributions of *n* emitted by the binary source approach that of a constitutive gene as *p*_*α*_ → 1.

## 3 Discussion

This manuscript shows a functional role for the unavoidable biochemical stochasticity observed inside the cells. We demonstrate that under proper regulation expression bursting of the transcription factors produced from a source gene maximizes the information transmitted to a receptor. We consider that the information to be exchanged between the genes is their promoter’s states since those are major determinants of gene expression levels. That is important for generating the internal conditions underpinning cell fate determination as observed during development, in which patterns of gene expression having striking spatio-temporal precision are observed.

The results presented at the Subsec. 2.1 show that one may increase the expression bursting of a binary gene to generate messages having arbitrarily small entropy. The reduction on the entropy and its maximal value depends both on the message’s bursting noise and on the probabilities governing the promoter states. We complement that analysis by computing the mutual information between the message and the promoter state. We verify that it has a maximum when the probability for the promoter to be ON is 1/2 and the bursting noise is approximately the average number of gene products. Therefore, we also show that, besides promoter state probabilities, the bursting of gene expression must also be regulated for the maximization of the information content of the message sent by the binary source.

Eq. 1 shows that bursting can be regulated by three distinct parameters. The value of *N* is determined by the capacity of the cell of synthesizing transcription factors while *p*_*α*_ is given by the enhancer’s state. Therefore, minimizing *ϵ* may be a useful complementary method for increasing level of bursting in the binary gene. Indeed, Subsec. 2.2 shows that we have reduced entropy and maximal mutual information for the slow switching genes. As expected, a very fast switching gene will generate a message having entropy approaching that of the constitutive gene having the same average expression levels.

The graphs of Subsec. 2.3 show that at maximal mutual information and reduced entropy, the distributions governing the message are bimodal. That implies on a slow switching and average duration of the ON or OFF states of the promoter being large enough for the number of transcription factors to reach steady state before the next switching. Hence, if we consider the conditional entropy given the promoter being ON, it will behave as a constitutive source. In other words, that regime permits the transcription factors to be a read out of the state of the promoter of the gene. As the parameter *ϵ* increases, the super Poissonian distributions change their shapes, *e*.*g*., becoming uniform-like, negative-binomial like, or Poissonian-like. Those regimes will have decreasing maximal mutual information and reduced quality on the information content of the message.

Optimization methods have been employed previously to verify that the bimodal distributions governing the signal, here interpreted as the number of transcription factors, enables maximal mutual information between the source and the receptor’s response [27, 40, 41]. That is done using a parameter-free approach useful for data analysis but not necessarily straightforward to be interpreted biologically and mechanistically. We use an approach based on an exactly solvable model with biologically interpretable, or observable, parameters. Hence, it can provide some guidance to synthetic biologists for designing gene circuits behaving as reliable (or unreliable) messaging devices. Additionally, it is a basic building block that can help on further understanding noise related biological phenomena [42, 43], *e*.*g*., occurring during development [44] or carcinogenesis [45, 46], in which information transmission plays a role.

The existence of a range of bursting regime for which mutual information is significant suggests that bursting is regulated in gene networks in which the promoter states are the information being exchanged between the source and receptor genes. There have been long term characterization of the enhancers as the gene structures responsible for regulating the promoter exposure for transcription and, consequently, average levels of gene products. Since gene expression bursting has also a functional role, one may wonder whether there exist a DNA structure surrounding a promoter that regulates its switching rates and, consequently, its bursting regimes, such that the gene network reliability is increased in contexts like development. Hence, it would be interesting to investigate experimentally the regulation of the promoter switching rates for multiple genes. Additionally, one might test the mutual information between a transcription factor and its target protein, or their promoters. Another interesting question is to test the information transfer between an external signal and genes involved in its processing. Our results also show bursty, non-constitutive promoters, have important biological role as elements of gene networks in which the important information is the promoter state. That contributes for the classification of noise function in gene expression besides self-repression as the mechanism responsible for noise reduction or homeostasis [47–50].

## 4 Materials and Methods

Here we assume that both the constitutive and the binary source are at the steady state regime. The source is interpreted as the gene encoding a transcription factor and the state of the source is determined as the promoter being ON or OFF. The constitutive source is always ON and the binary source switches at random between ON and OFF states. The regulation of a target gene depends on the interaction of the transcription factors with its enhancer. The binding of the transcription factors to an enhancer can be computed effectively as a function of the amounts of transcription factors around the DNA. Hence, the message emitted by the source is the number of transcription factors that are available for interaction with the enhancer of the receptor gene. The receptor gene promoter state is then determined by the action of the message on it, as either repressing or inducing transcription. The number of transcription factors denoted by *n* is a random variable and the noise on the message can be computed by the Fano factor, defined as the ratio between the variance of *n* to its average value. We denote the Fano factor by ℱ such that:

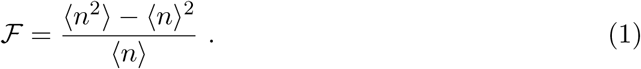

The Fano factor is a measure of how similar to the Poissonian a probability distribution is. The Poisson distribution has ℱ = 1 while ℱ *<* 1 characterizes a sub-Poisson distribution. A super-Poissonian distribution has ℱ > 1.

### 4.1 A stochastic model for a constitutive gene

We model a constitutive gene as a Poissonian birth and death process. The random variable *n* indicates the number of transcription factors available for regulating a receptor gene. The probability of finding *n* transcription factors is indicated by *ϕ*_*n*_. The synthesis and degradation rates are denoted by *k* and *ρ*, respectively. The probability *ϕ*_*n*_ is given by

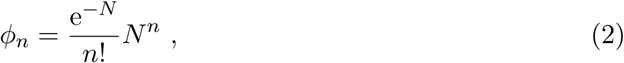

where

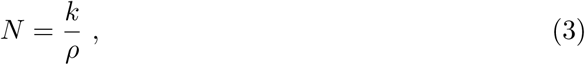

is the average number of transcription factors produced from a constitutive gene at the steady state regime. The variance of the Poissonian process is also *N*.

### 4.2 The stochastic model for the binary gene

We model the binary gene as two Poissonian birth and death processes coupled by a random telegraph process. The random telegraph process describes the switching of the promoter between ON and OFF states. When the promoter is ON (or OFF) proteins are synthesized at a rate denoted by *k* (or 0). The proteins degradation rate is denoted by *ρ*. There are two random variables, (*m, n*), where *m* indicates the state of the promoter as ON or OFF and *n* indicates the number of proteins. The probability of finding the promoter ON (or OFF) and *n* proteins is indicated by *α*_*n*_ (or *β*_*n*_). Using the exact solutions [36–38] for this model we have:

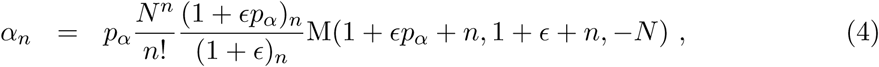

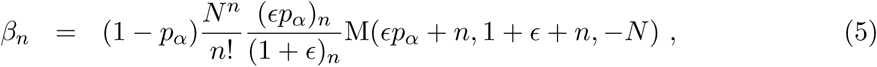

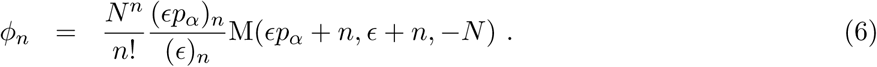

where the constants *p*_*α*_, *ϵ* and *N* are defined as:

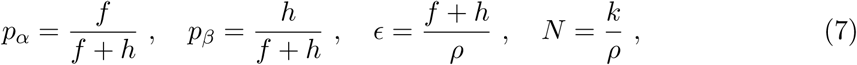

and *ϕ*_*n*_ indicates the marginal probability of finding *n* proteins, namely *ϕ*_*n*_ = *α*_*n*_ + *β*_*n*_.

M(*a, b, x*) denotes the KummerM function, defined as a power series:

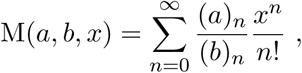

where the (*a*)_*n*_ = *a*(*a* + 1) … (*a* + *n* − 1) denotes the Pochhammer symbol [51].

### 4.3 Interpretation of the parameters *p*_*α*_, *ϵ*, and *N*

The parameters *p*_*α*_ and *p*_*β*_ is the steady state probability for the promoter to be at the ON and OFF state, with *p*_*α*_ + *p*_*β*_ = 1. *p*_*α*_ (or *p*_*β*_) give the fraction of a time interval *T* that the promoter will be ON (or OFF) while switching at random between its possible states.

The parameter *ϵ* is the ratio of the promoter switching rate to the transcription factors degradation rate. It can be interpreted as a ratio of the two time scales of the stochastic processes being coupled. For simplicity, we assume 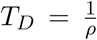 as the proteins lifetime, and 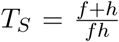 as the average time for the promoter to complete one switching cycle. Using the definitions at Eq. 7 we obtain

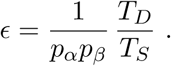

When *ϵ* ≪ 1 the switching cycle is long and proteins lifetime is shorter than the time spent by the promoter at ON or OFF state if *p*_*α*_ ∼ 1*/*2. When *ϵ* ≫ 1 the switching cycle is very short compared to the transcription factors lifetime and multiple switchings will happen while a transcription factor remains functional.

The parameter *N* is the average number of transcription factors if the binary gene remains ON all the time as if it was a constitutive gene.

### 4.4 The mean number and the conditional mean number of the stochastic model for a binary gene

The average number of transcription factors synthesized from the binary gene (⟨*n*⟩), and the average number of transcription factors given that the promoter is ON (denoted by ⟨*n*_*α*_⟩) are

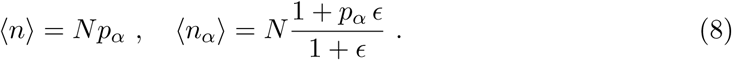

The mean number of transcription factors of the binary gene are dependent on the proportion of time that the promoter of the gene is ON. That indicates that the average amount of transcription factors will range from zero to *N* accordingly with *p*_*α*_. The conditional mean number of transcription factors given the promoter being ON is obtained from 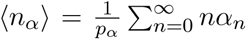. For very small values of *ϵ*, ⟨*n*_*α*_⟩ approaches *N*, which is the average for a Poissonian process. For large values of *ϵ* we have ⟨*n*_*α*_⟩ ∼ ⟨*n*⟩.

### 4.5 The variance and bursting noise of *n* on the stochastic model for a binary gene

The variance on *n*, denoted by *σ*^2^, and the Fano factor (ℱ) are, respectively,

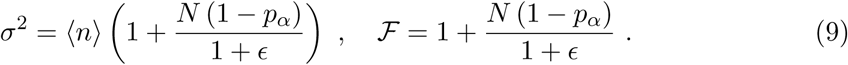

We interpret the variance and the Fano factor of *n* synthesized from a binary gene in comparison to that of a constitutive gene. Since the analysis is similar for both quantities, we focus on the Fano factor of *n* produced by the binary source is greater than that of the constitutive gene by a quantity *N* (1 − *p*_*α*_)*/*(1 + *ϵ*). This increment on ℱ is caused by the promoter switching. For *p*_*α*_ = 1 the promoter is fully ON, the binary source becomes constitutive and the increment on ℱ is null. For *p*_*α*_ = 0 we have a fully repressed gene. Additionally, when *ϵ* approaches infinity the bursting noise on the message sent by a binary source gene is similar to that of a constitutive gene for finite values of *N* and *p*_*α*_. For fixed values of *N* and *p*_*α*_ we have maximal bursting for *ϵ* approaching zero.

### 4.6 Entropy for the constitutive source

The Shannon’s entropy of a probability distribution *ϕ*_*n*_ is denoted by *H*(*n*) and defined as 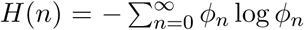, with the logarithm taken to base 2. For a Poissonian distribution having average value *N* the entropy is given by

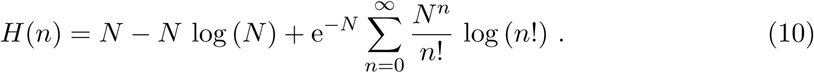

The entropy of the constitutive gene will be considered here as a reference value to be compared with that of the binary one.

### 4.7 Entropy, conditional entropy and mutual information for the binary source

Since it is not trivial to find a closed form to compute the logarithm of a KummerM function, we obtained the entropy and mutual information for the binary gene by computing their formulas numerically. The entropy of the message sent by the binary gene is:

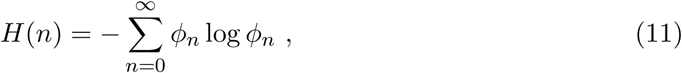

where *ϕ*_*n*_ is given by Eq. 6.

The conditional entropy of *n* given the promoter being ON or OFF is denoted by *H*_*α*_(*n*) or *H*_*β*_(*n*), respectively, and they are written as:

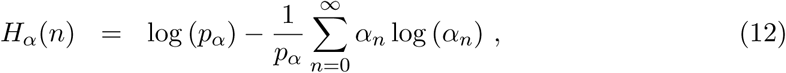

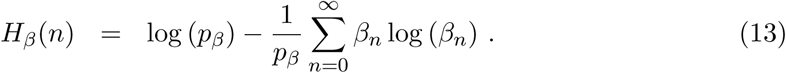

These formulas are obtained from the definition of the conditional entropy of *X* given 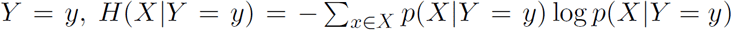, where we assume *X* being the number of transcription factors and *Y* the promoter state. Here *α*_*n*_, *β*_*n*_, and (*p*_*α*_, *p*_*β*_) are, respectively, given by Eq. 4, Eq. 5, and Eq. 7.

The mutual information between the message *n* and the source state being ON or OFF is given by

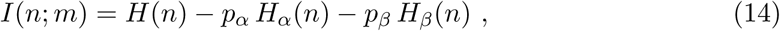

where *H*(*n*), *H*_*α*_(*n*), and *H*_*β*_(*n*) are, respectively, given by Eq. 11, Eq. 12, and Eq. 13.

## Author Contributions

Formal analysis, LRG, GG, GB, and AFR; Investigation, LRG, GG, and AFR; Visualization, LRG, GG, and AFR; Writing - original draft, AFR; Writing - review & editing LRG, GG, GB, and AFR; Funding acquisition, AFR.

## Acknowledgments

AFR is thankful to Yuri Suhov for introduction to Information Theory. We are thankful to Alan U. Sabino for useful comments.

